# Structured flexibility in recurrent neural networks via neuromodulation

**DOI:** 10.1101/2024.07.26.605315

**Authors:** Julia C. Costacurta, Shaunak Bhandarkar, David M. Zoltowski, Scott W. Linderman

## Abstract

The goal of theoretical neuroscience is to develop models that help us better understand biological intelligence. Such models range broadly in complexity and biological detail. For example, task-optimized recurrent neural networks (RNNs) have generated hypotheses about how the brain may perform various computations, but these models typically assume a fixed weight matrix representing the synaptic connectivity between neurons. From decades of neuroscience research, we know that synaptic weights are constantly changing, controlled in part by chemicals such as neuromodulators. In this work we explore the computational implications of synaptic gain scaling, a form of neuromodulation, using task-optimized low-rank RNNs. In our neuromodulated RNN (NM-RNN) model, a neuromodulatory subnetwork outputs a low-dimensional neuromodulatory signal that dynamically scales the low-rank recurrent weights of an output-generating RNN. In empirical experiments, we find that the structured flexibility in the NM-RNN allows it to both train and generalize with a higher degree of accuracy than low-rank RNNs on a set of canonical tasks. Additionally, via theoretical analyses we show how neuromodulatory gain scaling endows networks with gating mechanisms commonly found in artificial RNNs. We end by analyzing the low-rank dynamics of trained NM-RNNs, to show how task computations are distributed.

## 1 Introduction

Humans and animals show an innate ability to adapt and generalize their behavior across various environments and contexts. This suggests that the neural computations producing these behaviors must have flexible dynamics that are able to adjust to these novel conditions. Given the popularity of recurrent neural networks (RNNs) in studying such neural computations, a key question is whether this flexibility is (1) adequately and (2) accurately portrayed in RNN models of computation.

Traditional RNN models have fixed input, recurrent, and output weight matrices. Thus, the only way for an input to impact a dynamical computation is via the static input weight matrix. Prior work has shown that flexible, modular computation is possible with these models [1], but neurobiology suggests alternative mechanisms that may be at play. In particular, experimental neuroscience research has shown that synaptic strengths in the brain (akin to weight matrix entries in RNNs) are constantly changing — in part due to the influence of neuromodulatory signals [2].

Neuromodulatory signals are powerful and prevalent influences on neural activity and subsequent behavior. Dopamine, a well-known example, is implicated in motor deficits resulting from Parkinson’s disease and has been the subject of extensive study by neuroscientists [3]. For computational study, neuromodulators are especially interesting because of their effects on synaptic connections and learning [4]. In particular, neuromodulators have been shown to alter synaptic strength between neurons, effectively reconfiguring circuit dynamics [5].

In this work, we seek to incorporate a neuromodulatory signal into task-trained RNN models. We propose a model consisting of a pair of RNNs: a small “neuromodulatory” RNN and a larger, low-rank “output-generating” RNN. The neuromodulatory RNN controls the weights of the output-generating RNN by scaling each rank-1 component of its recurrent weight matrix. This allows the network to produce flexible, yet structured dynamics that unfold over the course of a task. We first review background work in both the machine learning and computational neuroscience literature. Next, we introduce the model and provide some intuition for the impact of neuromodulation on the model’s dynamics, relating it to the canonical long short-term memory (LSTM) network [6]. We end by presenting the performance and generalization capabilities of neuromodulated RNNs on a variety of neuroscience and machine learning tasks, showcasing the ability of a relatively simple, biologically-motivated augmentation to enhance the capacity of RNN models.

## 2 Background

First, we review related work in the theoretical neuroscience and machine learning literature.

### 2.1 Modeling neuromodulatory signals

Our work builds on a body of literature dating back to the 1980s, when pioneering computational neuroscientists added neuromodulation to small biophysical circuit models (for reviews, see [7–9]). These models consist of coupled differential equations whose biophysical parameters are carefully specified to simulate biologically-accurate spiking activity. As Marder [5] relates in her retrospective review, such neuromodulatory models were created in response to neuronal circuit models that viewed circuit dynamics as “hard-wired”. Neuromodulatory mechanisms offered an answer to experimental observations that anatomically fixed biological circuits were capable of producing variable outputs [10, 11]. Of particular relevance to this work, Abbott [12] showed in his 1990 paper that adding a neuromodulatory parameter to an ODE model of spiking activity allows a network of neurons to display capacity for both long- and short-term memory and gate the learning process, anticipating the LSTMs that would become prominent a few years later. We also draw comparisons between our model and the LSTM in the sections that follow. However, our work does not aim to model any specific biophysical system; rather, it aims to bridge the gap between these highly biologically-accurate models and general network models (i.e. RNNs) of neuronal activity by adding a biologically-motivated form of structured flexibility.

More recent attempts to model neuromodulation have taken advantage of increased computational power. Kennedy et al. [13] modeled modulatory influence in spiking RNNs by linearly scaling the firing rates of a subset of neurons. Papadopoulos et al. [14] also use spiking RNNs, and incorporate arousal-mediated modulatory signals to induce phase transitions. Stroud et al. [15] use a balanced excitatory/inhibitory RNN to model motor cortex, and incorporate modulatory signals as constant multipliers on each neuron’s activity. Liu et al. [16] study how neuromodulatory signals facilitate credit assignment during learning in spiking neural networks. Our work also aims to incorporate a modulatory signal into an existing neural network model, however, we specifically examine the potential of neuromodulation to augment task-trained low-rank RNNs.

The work of Tsuda et al. [17] is most similar to what we present here. The authors train RNNs using constant, multiplicative neuromodulatory signals applied to pre-specified subsets of the re-current weights. They show that these neuromodulatory signals allow an otherwise fixed network to perform variations of a task. In contrast, we employ time-dependent neuromodulatory signals that allow dynamics to evolve throughout tasks. Instead of pre-specifying the values and regions of impact of neuromodulators, we allow the model to learn the time-varying neuromodulatory signal and what segment of the neuronal population it impacts.

### 2.2 Hypernetworks

Our approach is closely related to recent work using hypernetworks to enhance model capacity in machine learning. Ha et al. [18] use small networks (termed hypernetworks) to generate parameters for layers of larger networks. In their HyperRNN, a hypernetwork generates the weight matrix of an RNN as the linear combination of a learned set of matrices. We also allow our neuromodulatory network to specify the weight matrix of a larger RNN as a linear combination of a learned set of matrices; however, our learned matrices are rank-1 to facilitate easier dynamical analysis and faster training. It is also worth noting that in practice, Ha et al. [18] simplify their HyperRNN so that the hypernetwork scales the rows of a learned weight matrix, which could be seen as postsynaptic scaling in our model. Similarly, von Oswald et al. [19] study the ability of hypernetworks to learn in the multitask and continual learning setting. They find that hypernetworks trained to produce task-specific weight realizations achieve high performance on continual learning benchmarks. In exploring potential neuroscience applications of their work, they remark that while their approach might be unrealistic, a hypernetwork that outputs lower-dimensional modulatory signals could assist in implementing task-specific mode-switching. We seek to obtain similar performance gains with a more biologically plausible model of neuromodulation.

### 2.3 Low-rank recurrent neural networks

For a variety of tasks of interest, measured neural recordings are often well-described by a set of lower dimensional latent variables [20, 21] (although alternative views have been presented [22, 23]). Likewise, artificial neural networks trained to solve tasks that mimic those found in neural experiments also often exhibit low-rank structure. Based on these findings, recurrent neural networks with low-rank weight matrices (also called low-rank RNNs, LR-RNNs) have emerged as a popular class of models for studying neural dynamics [24–26]. Importantly, low-rank RNNs are able to model nonlinear dynamics that evolve in a low-rank subspace, offering potential for visualization and interpretation. Here, we leverage low-rank RNNs as ideal candidates for neuromodulation, with each factor of the low-rank recurrence matrix becoming a possible target for synaptic scaling.

## 3 Neuromodulated recurrent neural networks

Motivated by the appealing dynamical structure of low-rank RNNs and the ability of neuromodulation to add structured flexibility, we propose the *neuromodulated RNN* (NM-RNN). The NM-RNN consists of two linked subnetworks corresponding to neuromodulation and output generation. The output generating subnetwork is a low-rank RNN, which admits a natural way to implement neuromodulation. We allow the output of the neuromodulatory subnetwork to scale the low-rank factors of the output generating subnetwork’s weight matrix. In particular, we propose incorporating neuromodulatory drive via a coupled ODE with neuromodulatory subnetwork state ***z***(*t*) ∈ ℝ^*M*^ and output-generating state ***x*** (*t*) ∈ ℝ^*N*^ :

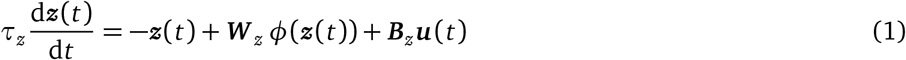

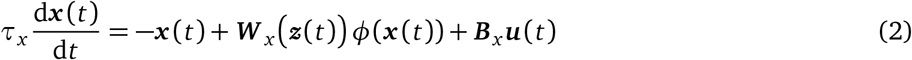

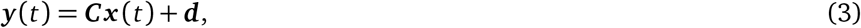

where the dynamics matrix ***W*** _*x*_ (***z***(*t*)) is a function of the neuromodulatory subnetwork state ***z***(*t*) via a neuromodulatory signal ***s*** (***z***(*t*)) ∈ ℝ^*K*^, which scales each low-rank component of ***W*** _*x*_ :

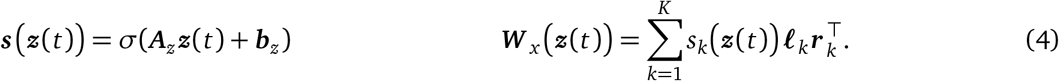

The output-generating subnetwork is modeled as a size-*N* low-rank RNN where ***u***(*t*) ∈ ℝ^*P*^ are the inputs, ***B***_*x*_ ∈ ℝ^*N×P*^ are the input weights, and *φ*(·) is the tanh nonlinearity. The neuromodulatory subnetwork is modeled as a small “vanilla” RNN with its own time-constant *τ*_*z*_, recurrence weights ***W***_*z*_ ∈ ℝ^*M×M*^, and input weights ***B***_*z*_ ∈ ℝ^*M×P*^. To limit the capacity of the neuromodulatory subnetwork, we take its dimension *M* to be smaller than the output-generating subnetwork’s dimension *N*. We take *τ*_*z*_ ≫ *τ*_*x*_ since neuromodulatory signals are believed to evolve relatively slowly [27]. The neuromodulatory subnetwork state ***z***(*t*) alters the rank-*K* dynamics matrix ***W***_*x*_ ∈ ℝ^*N×N*^ via a *K*-dimensional linear readout ***s*** (***z***(*t*)). The components of ***s*** act as linear scaling factors on each rank-1 component 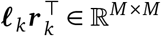 of ***W***_*x*_. For ease of notation, in the rest of the text we write ***s*** (*t*) to mean ***s*** (***z***(*t*)). The output ***y*** (*t*) ∈ ℝ^*O*^ of the NM-RNN is a linear readout of ***x*** (*t*).

This augmentation to the RNN framework allows for structured flexibility in computation. In traditional RNNs, the recurrent weight matrix is fixed, and thus the inputs to the system can only perturb the state of the network. In the NM-RNN, the neuromodulatory subnetwork can use information from the inputs to dynamically up- and down-weight individual low-rank components of the recurrent weight matrix, offering greater computational flexibility. As we will see below, this flexibility also allows the network to reuse dynamical components across different tasks and task conditions.

### 3.1 Mathematical intuition

To gain some intuition for the potential impacts of neuromodulation on RNN dynamics, first consider the case where ***W*** _*x*_ is symmetric (i.e., ***𝓁***_*k*_ = ***r*** _*k*_ ∀*k*) with the {***𝓁***_*k*_}_1≤*k*≤*K*_ forming an orthonormal set, where the nonlinearity is removed (i.e., *φ*(*x*) = *x*), and where there are no inputs (i.e., ***u***(*t*) = 0 ∀*t*). We can then reparameterize the system with a new hidden state ***w*** (*t*) = ***L***^⊤^ ***x*** (*t*), where ***L*** ∈ ℝ^*N×K*^ is the matrix whose columns are ***𝓁***_*k*_ (so that ***L***^*T*^ ***L*** = **I**). This produces decoupled dynamics:

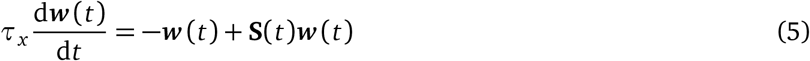

where ***S***(*t*) = diag (***s*** (*t*)). Solving this ODE gives an equation for the components of ***w*** (*t*):

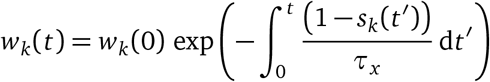

From this equation and the visualization in fig. 2A, we see that the decay rate of each component *w*_*k*_(*t*) is governed by its corresponding neuromodulatory signal *s*_*k*_(*t*). In this way, ***s*** (*t*) can effectively speed up or slow down decay of dynamic modes, similar to gating in an LSTM.

**Figure 1:**
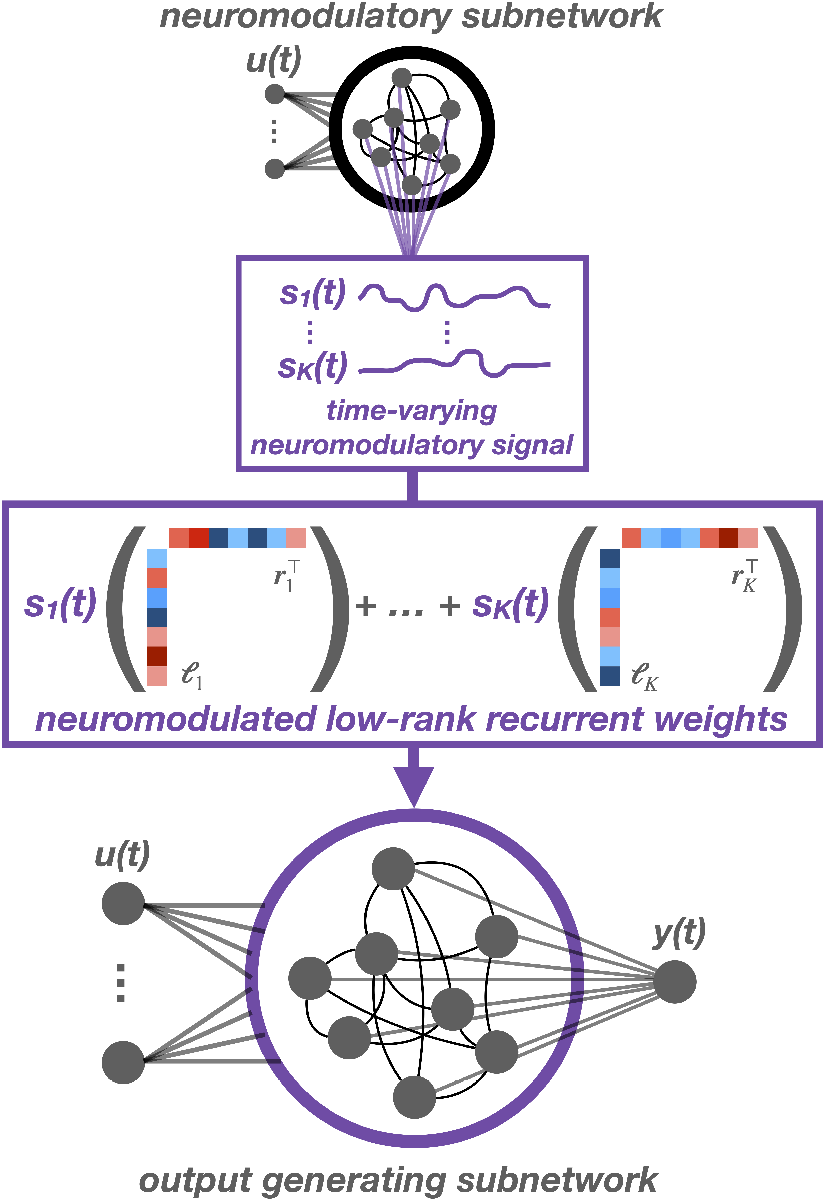
The NM-RNN consists of two subnetworks: a low-rank subnetwork that generates the output (bottom) and a smaller, full-rank neuromodulatory subnetwork (top).

**Figure 2:**
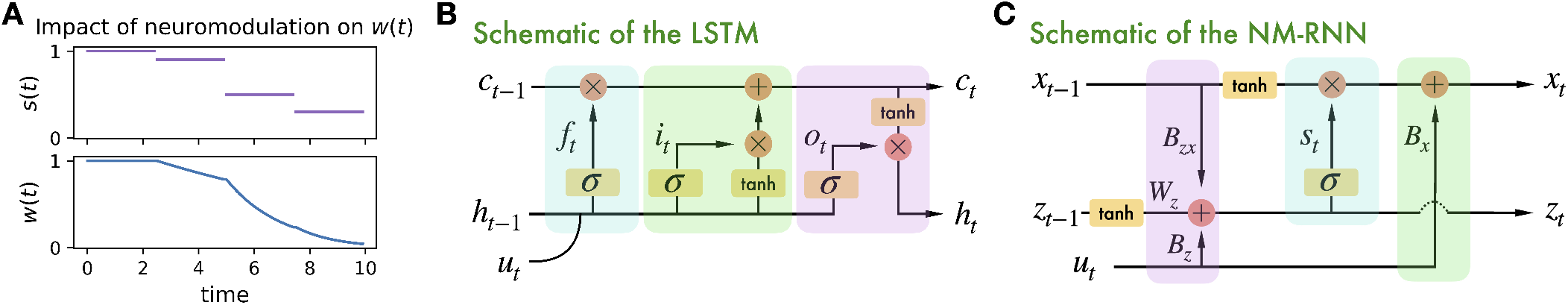
**A**. Illustration of how neuromodulatory signals *s*(*t*) affect decay rates of the state *w*(*t*) in a 1-D, simplified model. **B. & C**. Visual comparison of an LSTM and an NM-RNN. Corresponding parts of the networks are highlighted with shaded rectangles. Blue: a forget gate computation. Green: an input gate to recurrent dynamics. Purple: recurrent feedback onto the modulatory state variable.

### 3.2 Connection to LSTMs

Having observed that neuromodulation can alter the timescales of dynamics, note further that the low-rank update for the linearized NM-RNN in eq. (5) mirrors the cell-state update equation for a long short-term memory (LSTM) cell [6]. Specifically, the neuromodulatory signal ***s*** (*t*) resembles the forget gate of an LSTM (fig. 2B). Indeed, as in eq. (5), if we linearize the output-generating subnetwork of the NM-RNN and assume that ***L*** = ***R*** (so that ***W*** _*x*_ is symmetric), and ***L***^*T*^ ***L*** = *I*, then for ***w*** (*t*) = ***L***^*T*^ ***x*** (*t*) and *τ*_*x*_ = 1, the discretized low-rank dynamics are given by

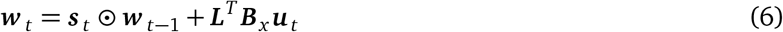

LSTMs have two states that recurrently update across each timestep *t*: a hidden state 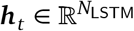 and a cell state 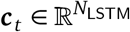. Equation (6) closely mirrors the cell-state update of the LSTM:

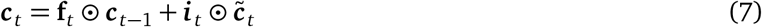

Here, the forget gate **f**_*t*_ is a form of modulation that depends on the LSTM hidden state ***h***_*t*_, much like the NM-RNN’s neuromodulatory signal ***s*** (*t*) is a form of modulation depending on the NM-RNN’s neuromodulatory subnetwork state ***z***(*t*). The second term 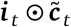 can be viewed as a gated transformation of the input signal ***u***(*t*). In fact, under suitable assumptions, we show that the dynamics of an NM-RNN can be reproduced by those of an LSTM (see Theorem 1).

As a gated analog of the RNN, the LSTM has enjoyed greater success than ordinary RNNs in performing tasks that involve keeping track of long-distance dependencies in the input signal [28]. Thus, highlighting the connection between the NM-RNN and LSTM suggests NM-RNNs may be able to model long-timescale dependencies better than regular RNNs (see Section 6).

## 4 Time interval reproduction

To evaluate the potential of neuromodulated RNNs to add structured flexibility, we first consider a timing task since neuromodulators such as dopamine are implicated in time measurement and perception [29]. In the Measure-Wait-Go (MWG) task (fig. 3A) [30], the network receives a 3-channel input containing the measure, wait, and go cues. The network must measure the interval between the measure and wait cues and reproduce it at the go cue by outputting a linear ramp of the same duration. Tasks such as this one are commonly used to study the neural underpinnings of timing perception in humans and non-human primates [31, 32].

**Figure 3:**
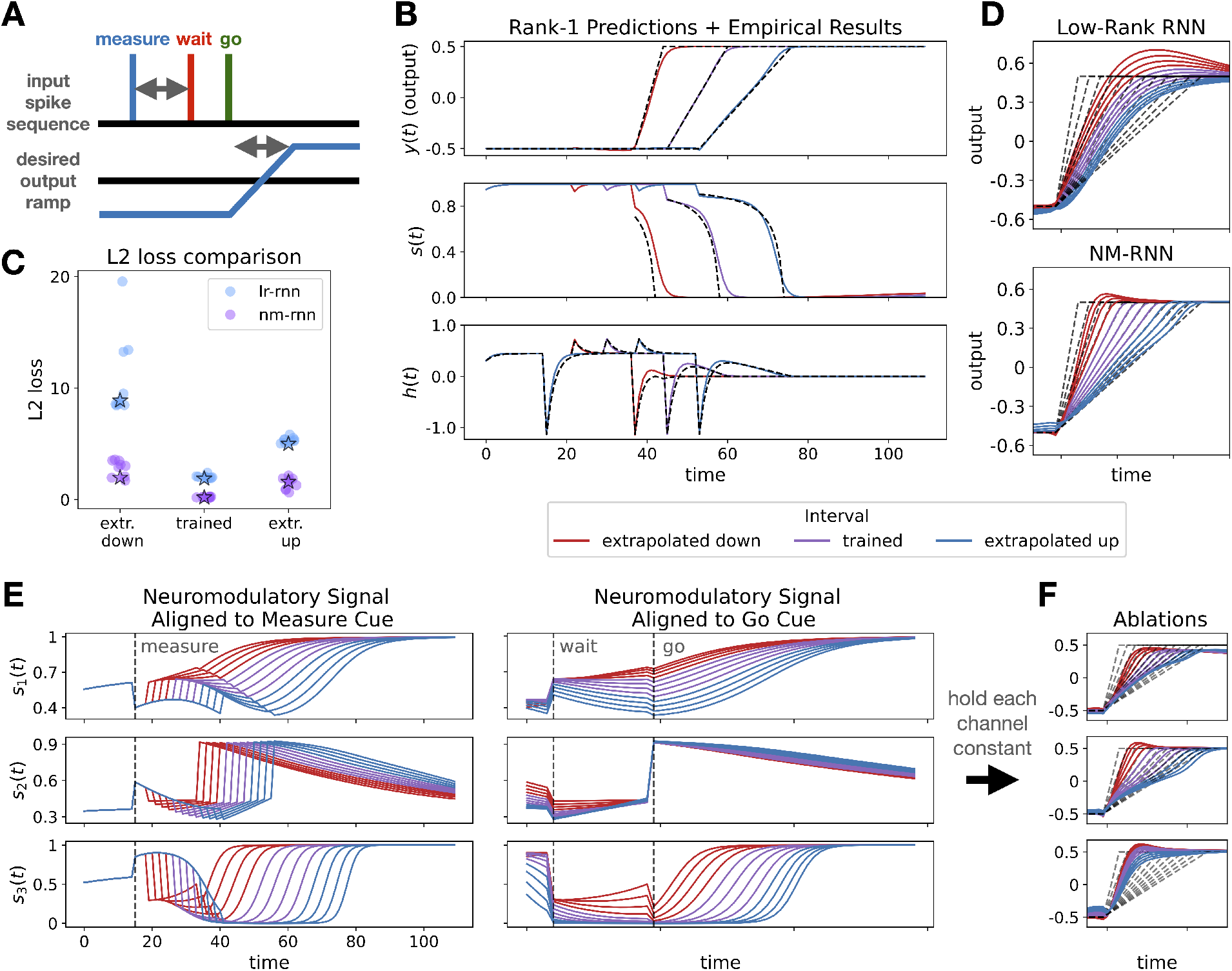
**A**. Visualization of Measure-Wait-Go task. **B**. Theoretical predictions (dashed lines) match closely with empirical results (solid lines) for rank-1 network. **C**. L2 loss comparsion for parameter-matched LR-RNNs and NM-RNNs. Performance of visualized models are starred. **D**. Comparison of model-generated output ramps for both trained and extrapolated intervals. **E**.Three-dimensional neuromodulatory signal *s* (*t*) for trained/extrapolated intervals. Left, traces are aligned to start of trial. Right, traces are aligned to ‘go’ cue. **F**. Resulting output traces when ablating each component of *s* (*t*). In all panels, colors reflect trained/extrapolated intervals (see legend). For output plots, dashed grey lines are targets. Additional model visualizations in figs. S1 and S2.

### 4.1 Experiment matches theory for rank-1 networks

To investigate how the NM-RNN’s neuromodulatory signal is constrained by the task requirements, we first consider a class of analytically tractable NM-RNNs: networks for which the output-generating subnetwork is linear (i.e., the tanh nonlinearity is replaced with the identity function) and rank-1. In this case, there is one pair of row and column factors, ***𝓁*** and ***r***, respectively. If the target output signal is given by *f* (*t*) and there are no inputs, then an NM-RNN that successfully produces this output signal will precisely have the neuromodulatory signal,

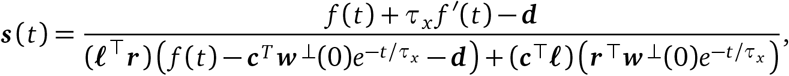

where our readout is *y* = ***c***^⊤^ ***x*** + ***d*** and 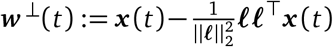 is the component of ***x*** (*t*) evolving outside of the column space of ***𝓁*** (viewed as a matrix in ℝ^*N*×1^). For the full derivation, see Appendix B. If we assume further that ***w*** ^⊥^(0) is sufficiently small and *τ*_*x*_ is also sufficiently small, then we may make the approximation,

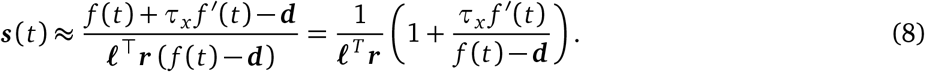

In the MWG task, no inputs are provided to the network during the ramping period following the go cue, so eq. (8) applies. In fig. 3B, we show that the neuromodulatory signal of a trained rank-1 NM-RNN during the ramping period matches closely with the theoretical prediction made by eq. (8) for both trained and extrapolated target intervals.

### 4.2 Improved generalization and interpretability on timing task for rank-3 networks

To continue our analysis of NM-RNNs on the MWG task, we increase the rank of the output-generating subnetwork to three. We do this to compare to the networks shown in Beiran et al. [30] and to showcase networks with more degrees of structured flexibility. In Beiran et al. [30], the authors show that rank-3 low-rank RNNs perform and extrapolate better on this task when provided a tonic context-dependent input, which varies depending on the length of the desired interval. As we have mentioned, such sensory inputs to the network may only alter the resulting dynamics by being passed through the input weight matrix. We propose the NM-RNN as an alternative mechanism by which inputs may alter the network dynamics.

We trained parameter-matched LR-RNNs (*N* = 106, *τ* = 10) and NM-RNNs (*N* = 100, *M* = 5, *τ*_*x*_ = 10, *τ*_*z*_ = 100) to reproduce four intervals, then tested their extrapolation to longer and shorter intervals. In each of the networks, the low-rank matrix was chosen to have rank 3, as in Beiran et al. [30]. In fig. 3B, we plot the *L*_2_ losses for ten instances of each model, showing that the NM-RNN consistently achieves a lower loss on both the trained and extrapolated intervals. In fig. 3C, we then show outputs for two typical networks (performance of these visualized networks indicated by stars in fig. 3B). The outputs of the NM-RNN have more accurate slope and shape for both trained and extrapolated intervals, with the LR-RNN struggling on shorter extrapolated intervals.

We then investigate how the neuromodulatory signal ***s*** (*t*) contributes to the task computation. Figure 3E shows the three dimensions of ***s*** (*t*) plotted over the full range of trained and extrapolated intervals. Each dimension shows activity correlated to particular stages of the task. In fig. 3E (right), we see that *s*_1_(*t*) and *s*_3_(*t*) have activity highly correlated to the measured interval. In particular, after the go cue, *s*_3_(*t*) ramps at different speeds to saturate at 1, depending on the length of the interval. Figure 3F shows the result of ablating each dimension of ***s*** (*t*) by keeping that component fixed around its initial value. We see that performance suffers in all cases, especially when ablating the effect of *s*_1_(*t*) and *s*_3_(*t*). Most dramatically, ablating *s*_3_(*t*) destroys the ability of the network to change the slope of the output ramp appropriately. These results show how the network uses its neuromodulatory signal to generalize across task conditions.

## 5 Reusing dynamics for multitask learning

Next, we move beyond generalization within a single task to investigate the capabilites of the NM-RNN when switching between tasks. There has been recent interest in studying how neural network models might reassemble learned dynamical motifs to accomplish multiple tasks [1, 33]. Driscoll et al. [1] showed that an RNN trained to perform an array of tasks shares modular dynamical motifs across task periods and between tasks. With this result in mind, we were curious how the NM-RNN might use its neuromodulatory signal to flexibly reconfigure dynamics across tasks.

We performed our analysis using the four-task set from Duncker et al. [34], which includes the tasks DelayPro, DelayAnti, MemoryPro, and MemoryAnti illustrated in fig. 4A. In the DelayPro task, the network receives a three-channel input consisting of a fixation input and two sensory inputs which encode an angle *θ* ∈ [0, 2*π*) as (sin(*θ*), cos(*θ*)). The fixation input starts and remains at 1, then drops to 0 to signal the start of the *readout* period, when the network must generate its response. The sensory inputs appear after a delay, and persist throughout the trial. The goal of the network is to produce a three-channel output which reproduces the fixation and sensory inputs. In the MemoryPro task, the sensory inputs disappear before the readout period, requiring the network to store *θ*. In the ‘Anti’ tasks, the networks must instead produce the opposite sensory outputs, (sin(*θ* + *π*), cos(*θ* + *π*)), during the readout period. The task context is fed in as an additional one-hot input. These tasks are analogous to variants of the center-out reaching task, which has been used to study the neural mechanisms of arm movement in non-human primates [35].

**Figure 4:**
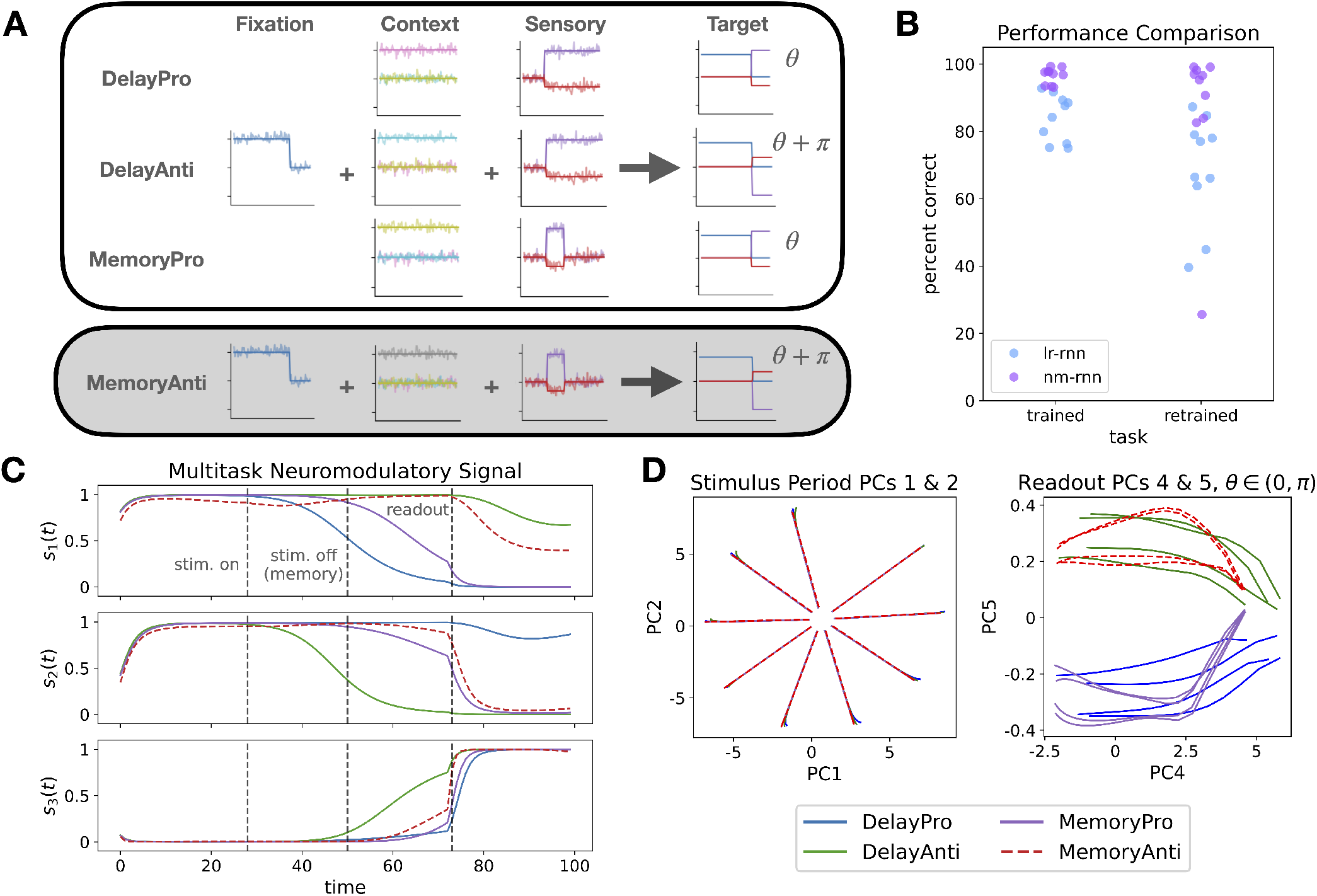
**A**. Depiction of inputs and targets for four tasks. Networks were trained on the first three tasks, then contextprocessing weights were retrained on the fourth task (in grey). **B**. Comparison of percent correct metric between parametermatched low-rank and neuromodulated RNNs, for initial three tasks and retrained MemoryAnti task. **C**. Neuromodulatory signal for an example network. **D**. Dynamical analysis of network activity during different stages of the tasks. (Left) PCs 1&2 during the stimulus period show a ring attractor which stores the measured angle. (Right) Sign of PC5 during readout period corresponds to Pro/Anti. Additional model visualizations in figs. S3 and S4.

To study the potential of NM-RNNs to flexibly reconfigure dynamics to perform a new task, we only fed the contextual inputs to the neuromodulatory subnetwork, and not to the output-generating subnetwork. This required the model to reuse the output-generating subnetwork’s weights when adding a new task. We trained an NM-RNN to perform the first three tasks in the set (DelayPro, DelayAnti, MemoryPro), then froze the weights of the output-generating subnetwork and retrained only the neuromodulatory subnetwork’s weights on the fourth task, MemoryAnti. We compared this to retraining the input weights of a LR-RNN, to investigate two strategies of processing context.

We trained parameter-matched LR-RNNs (*N* = 100, *τ* = 10) and NM-RNNs (*N* = 100, *M* = 20, *τ*_*x*_ = 10, *τ*_*z*_ = 100) in this training/retraining framework. Figure 4B shows performance of example networks on the trained and retrained tasks, using the percent correct metric from Driscoll et al. [1]. While both models are able to learn the first three tasks, the NM-RNN more consistently performs the fourth task with high accuracy. This performance gain is not the result of retraining more parameters; in fact, due to the contrasting sizes of the neuromodulatory and low-rank subnetworks, the input weight matrix of the comparison low-rank RNN contains more parameters than the entire neuromodulatory subnetwork, since it must process all inputs (context, sensory, and fixation). To see exactly how the recurrent dynamics were rearranged for this new task, we plotted the neuromodulatory signal of an example network for learned and extrapolated tasks in fig. 4C.

We then analyzed the dynamical structure of one of the NM-RNNs by performing PCA on the output-generating subnetwork’s state ***x*** (*t*) for a variety of input angles. Figure 4D (left) shows the first two PCs of the neural activity during the stimulus presentation period (before the stimulus shut off for Memory trials). During this period, the neural activity spreads out to arrange on a ring according to the measured angle. After the stimulus disappears in the MemoryPro/Anti tasks, the neural activity decays back along these axes, but it is still decodable based on its angle from the origin (see fig. S4). To find how this model encoded Pro/Anti versions of tasks, we performed another PCA on the neural activity during the readout period. As shown in fig. 4D (right), the sign of PC5 during this period is correlated with whether the task is Pro or Anti. Curiously, the positive/negative relationship flips for *θ* ∈ (*π*, 2*π*), likely relating to the symmetric structure of sine and cosine. These results show the ability of the NM-RNN to flexibly reconfigure the dynamics of the output-generating subnetwork, both to solve multiple tasks simultaneously and to generalize to a novel task.

## 6 Capturing long-term dependencies via neuromodulation

Inspired by the similarity between the coupled NM-RNN and the LSTM (see section 3.2), we designed a sequence-related task with long-term dependencies, called the *Element Finder Task* (EFT). On this task, gated models like the NM-RNN outperform ordinary RNNs. When endowed with suitable feedback coupling from the output-generating subnetwork to the neuromodulatory subnetwork, the NM-RNN demonstrates LSTM-like performance on the EFT, while vanilla RNNs (with matched parameter count) fail to solve this task.

In the EFT (fig. 5A), the input stream consists of a query index, *q* ∈ {0, 1, …, *T* − 1} followed by a sequence of *T* randomly chosen integers. The goal of the model is to recover the value of the element at index *q* from the sequence of integers. At each time for *t* ≥ 1, the *t*th element of the sequence is passed as a one-dimensional input to the model. At time *t* = *T*, the model must output the value of the element at index *q* in the sequence. For our results (shown below), we took *T* = 25.

**Figure 5:**
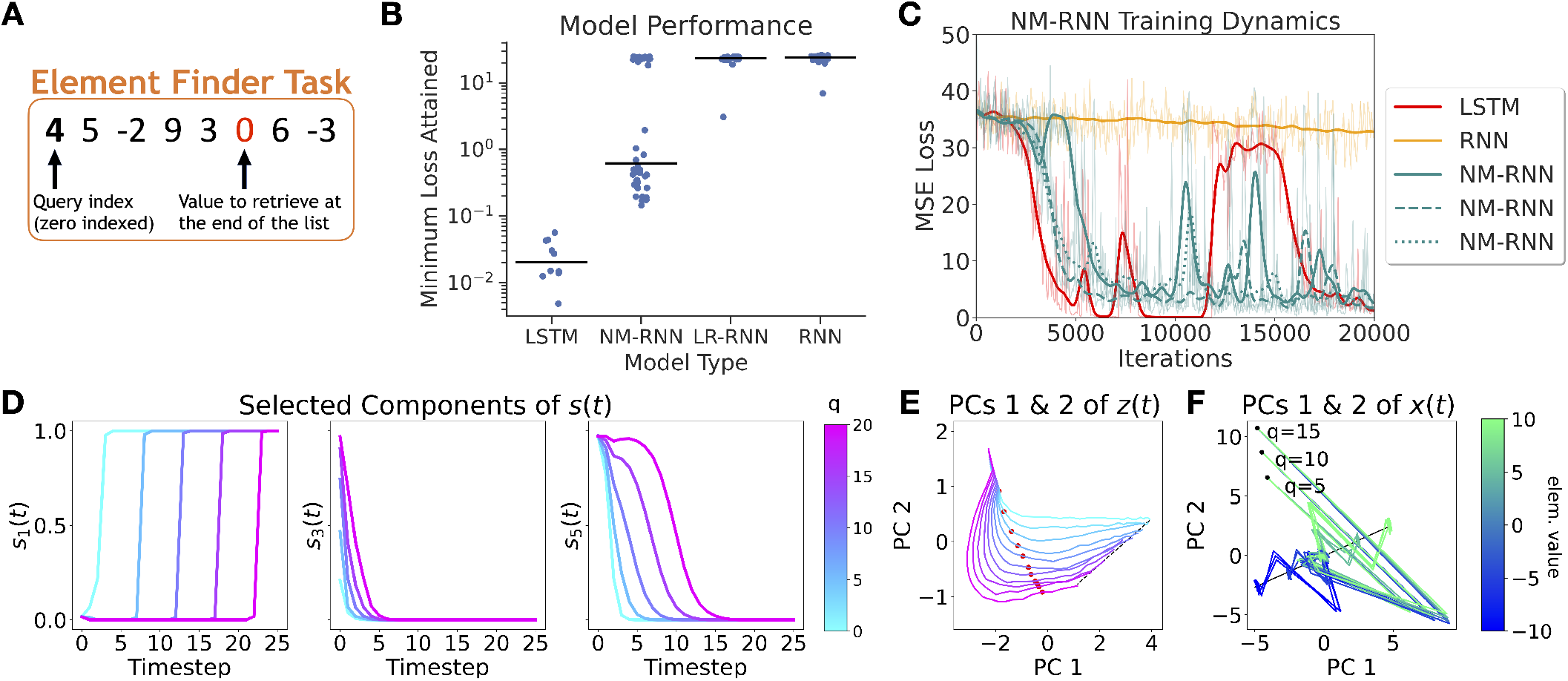
**A**. Visualization of the Element Finder Task. **B**. MSE losses attained across multiple runs in different classes of models trained on the EFT (median is indicated by black lines). **C**. Training loss curves for selected models. **D**. Visualization of selected components of *s* (*t*) for an example NM-RNN, shown across different query indices. **E**. Trajectories for the top two PCs of *z*(*t*) across different query indices. The different trajectories converge to an approximate line attractor (black) encoding query index. The time at which the queried element arrives is marked in red. **F**. Top two PCs of *x* (*t*), visualized for different query indices and target element values. Each trajectory converges to a fixed point on an approximate line attractor encoding element value. Each curve shown in **D, E**, and **F** is averaged over 100 trials. Additional visualizations are shown in figs. S5 and S6

We trained several NM-RNNs, LR-RNNs, full-rank RNNs, and LSTMs on the EFT, conserving the total parameter count across networks. To emphasize its connection to the LSTM, each NM-RNN included an additional feedback coupling from ***x*** (*t*) to ***z***(*t*):

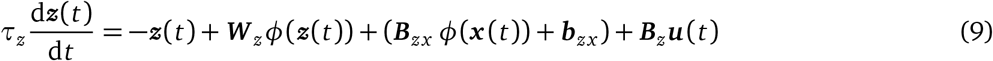

Each model used a linear readout with no readout bias. The resulting performances of each model tested are shown in fig. 5B. Figure 5C moreover illustrates the learning dynamics (as measured by MSE loss) for a single run of selected networks. Like LSTMs, NM-RNNs successfully perform the task, whereas lowand full-rank RNNs largely fail to do so.

To understand how a particular NM-RNN (*N* = 18, *M* = 5, *K* = 8, *τ*_*x*_ = 2, *τ*_*z*_ = 10) uses neuromodulatory gating to solve the EFT, we visualize the trial-averaged behavior of different components of ***s*** (*t*) across query indices (*q* = 5, 10, 15, 20), revealing that certain components of ***s*** (*t*) transition between 0 and 1 on a timescale correlated to the query index *q* (fig. 5D; left and right); while other components zero out (fig. 5D; middle). Visualizing a low-dimensional projection of ***z***(*t*) across different query indices reveals that ***z***(*t*) settles to a fixed point on an approximate line attractor encoding query index *q* (fig. 5E). These findings show that ***z***(*t*) attends to the query index, facilitating gate-switching behavior in ***s*** (*t*) upon arrival of the queried element.

Next, we analyze ***x*** (*t*) by visualizing its top 2 principal components across each combination of the query indices *q* = 5, 10, and 15 and the target element values −10, −5, 0, 5, and 10 (fig. 5F). Trajectories with different element values but the same query index start at the same location. Each trajectory converges towards the origin, and upon arrival of the query timestep, rapidly moves to a fixed point on an approximate line attractor that encodes element value. The arrangement of fixed points along this line moreover preserves the ordering of their corresponding element values. In summary, these results show that the NM-RNN solves the EFT by distributing its computations across the neuromodulatory subnetwork, which attends to the query index, and the output-generating subnetwork, which retrieves the target element value.

## 7 Discussion

As we have shown, neuromodulated RNNs display an increased ability to both perform and generalize on tasks, both in the single-task setting and between different tasks. This performance is enabled by the structured flexibility neuromodulation adds to the dynamics of the network. Curiously, the gating-like dynamics introduced by adding neuromodulation create strong comparsions (and even equivalence) to the canonical LSTM.

### Limitations

One limitation of this work relates to the scale of the networks tested. Our networks were on the scale of *N* ≈ 100 neurons at their largest, as opposed to other related works which use neuron counts in the thousands. However, we found that this number of neurons was adequate to perform the tasks we presented. We also have yet to compare our results to neural data, limiting our ability to draw biological conclusions.

### Future Work

We are excited at the potential of future work to further bridge the gap between biophysical and recurrent neural network models. To expand on the NM-RNN model, we aim to embrace the broad range of roles neuromodulation can play in neural circuits. Potential future avenues include: (1) sparsifying the rank-1 components of the recurrent weight matrix to better imitate the ability of neuromodulators to act on spatially localized subpopulations of cells; (2) changing the readout function of ***s*** (*t*) to enable it to take both negative and positive values, in line with the ability of neuromodulators to act in both excitatory and inhibitory manners; and (3) investigating how different neuromodulatory effects may act on different timescales, both during task completion and learning over longer timescales [4, 5]. More generally, each neuron (or synapse) could have an internal state beyond its firing rate which is manipulated by neuromodulators, as in recent work investigating the role of modulation in generating novel dynamical patterns [36, 37]. Beyond neuromodulators, there exist a multitude of extrasynaptic signaling mechanisms in the brain, such as neuropeptides and hormones, which could function similarly in some cases, but also suggest new modeling directions.

In this work, we only analyzed networks post-training. We are also curious how our computational mechanism of neuromodulation impacts the network during learning. Prior work has modeled the role of neuromodulation in learning, for example, by augmenting the traditional Hebbian learning rule with neuromodulation to implement a three-factor learning rule [38], and by using neuromodulation to create a more biologically plausible learning rule for RNNs [39]. An interesting direction for future work is to study whether the neuomodulatory signal in the NM-RNN could produce similar learning dynamics.

## Acknowledgements

We thank Laura Driscoll, Lea Duncker, Gyu Heo, Bernardo Sabatini, and the members of the Linderman Lab for helpful feedback throughout this project. This work was supported by grants from the NIH BRAIN Initiative (U19NS113201, R01NS131987, & RF1MH133778) and the NSF/NIH CRCNS Program (R01NS130789). J.C.C. is funded by the NSF Graduate Research Fellowship, Stanford Graduate Fellowship, and Stanford Diversifying

Academia, Recruiting Excellence (DARE) Fellowship. D.M.Z. is funded by the Wu Tsai Interdisciplinary Postdoctoral Research Fellowship. S.W.L. is supported by fellowships from the Simons Collaboration on the Global Brain, the Alfred P. Sloan Foundation, and the McKnight Foundation. The authors have no competing interests to declare.

## A Formal connection between NM-RNNs and LSTMs

In this section, we mathematically formalize the connection between the NM-RNN and the LSTM. We show that, when considering suitable *linearized* analogs of the NM-RNN and the LSTM, the internal states and outputs of an NM-RNN may be reproduced by an LSTM. To that end, we begin by stating the necessary definitions.

First, we define a *linearized NM-RNN* via the following discretized dynamics:

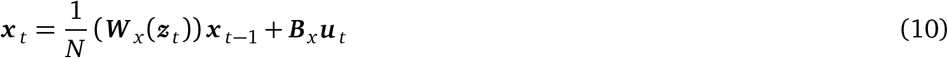

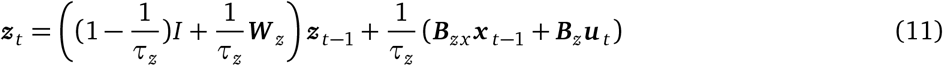

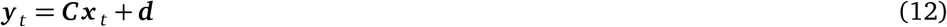

As defined in the main text, we have that ***W*** _*x*_ (***z*** _*t*_) = ***LS***_*t*_ ***R***^*T*^ ∈ ℝ^*N×N*^, where *S*_*t*_ = diag(***s*** _*t*_) and ***s*** _*t*_ = *σ*(***A***_*z*_ ***z*** _*t*_ + ***b***_*z*_) ∈ ℝ^*K*^. Note that we have added a 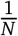 prefactor to ***W*** _*x*_ in eq. (10). However, this does not fundamentally change the computation in Equation eq. (10) because we may imagine that this prefactor is absorbed by the matrices ***L*** and ***R***. Explicitly having the 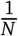 prefactor will be make the presentation of Proposition 1 more convenient.

Observe that the linearized NM-RNN is an NM-RNN with *τ*_*x*_ = 1 and where all nonlinear transfer functions have been made to be the identity function. Notably, we have also added a *feedback coupling* term (given by the feedback weights ***B***_*zx*_) from ***x*** (*t*) to ***z***(*t*); this will serve to enhance the connection between this model and the LSTM.

Turning to the LSTM, we similarly define a suitable linearized relaxation – the *semilinear LSTM* – given by the following equations:

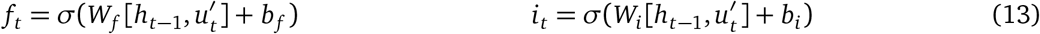

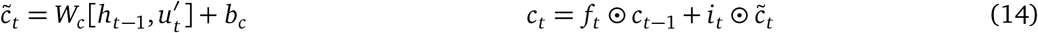

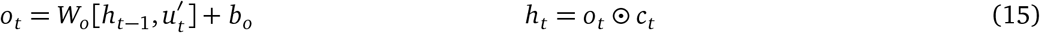

Here, *σ* denotes the sigmoid nonlinearity, 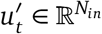 denotes the input, and 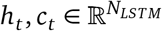 are the hidden and cell states, respectively. The notation [·, ·] signifies concatenation of vectors. Each of the parameters *W*_*k*_, *b*_*k*_, for *k* ∈ { *f, i, c, o*}, are learnable.

### A.1 Conditions leading to equivalence

Now, we formalize the connection between these two classes of models. It will be revealing to analyze the linearized NM-RNN in the case where ***L*** = ***R***.

**Proposition 1**. *Consider a rank-K linearized NM-RNN, where the hidden size of the (linearized) output-generating network is N, and the hidden size of the neuromodulatory RNN is M. Moreover, assume that* ***L*** = ***R*** *and that the columns of* 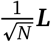 *are pairwise orthogonal, in the sense that* 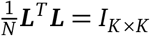. *Finally, across all input sequences* ***u*** _*t*_ *on which the NM-RNN is tested, assume that the components of the states* ***x*** _*t*_ *and* ***z*** _*t*_ *are uniformly bounded over all timesteps. Then, this model’s underlying K-dimensional dynamics (given by the low-rank variable* 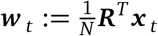*), as well as its neuromodulatory states* ***z*** _*t*_ *and outputs* ***y***_*t*_ ∈ ℝ ^*O*^, *can be reproduced within a semilinear LSTM with a linear readout, whose inputs are* 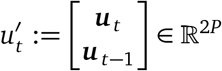. *Here*, ***u*** _*t*_ ∈ ℝ *denotes the input to the NM-RNN at time t (with* ***u***_−1_ := 0 ∈ ℝ^*P*^*by convention). Moreover, such a semilinear LSTM can be made to have hidden size K* + *M* + *P and utilize O* max{*K, M, PK, PM, OK, OP*} *learnable parameters*.

*Proof of Proposition 1*. Assuming ***x***_*t*_ has uniformly bounded components, there exists some constant *H* ∈ ℝ_+_ such that |(***x***_*t*_)_*i*_| *< H* for each *i* ∈ {1, …, *N*} and all *t*. Consequently, this means 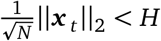 for all *t*. Now, consider the update equation for ***x*** _*t*_ in the linearized NM-RNN:

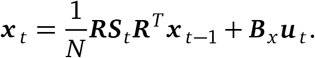

For ***R***_*i*_ the *i*th column of ***R***, the given orthogonality condition implies that 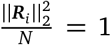,or 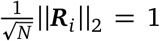 in the large *N* limit. Defining 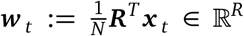 to the rank-*K* representation of ***x*** _*t*_ ∈ ℝ^*N*^, we have that 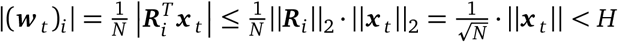. That is, the low-rank mode corresponding to ***x*** _*t*_ also has uniformly bounded components over time, i.e., the underlying *K*-dimensional dynamics of the output-generating network do not diverge.

Now, to show that the dynamics of a linearized NM-RNN satisfying the conditions given in the theorem statement can be replicated by a semilinear LSTM, we must effectively show that each of the update equations for the linearized NM-RNN (i.e., eqs. (10), (11) and (12)) can be suitably replicated through the semilinear LSTM architecture. In essence, we must suitably map the parameters of the given NM-RNN onto those of a semilinear LSTM.

First, we analyze the update equation for ***w*** _*t*_ in the output-generating network. Its corresponding *K*-dimensional (low-rank) update is

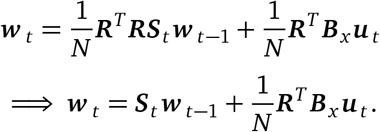

Accordingly, define the cell state *c*_*t*_ of the corresponding semilinear LSTM to be

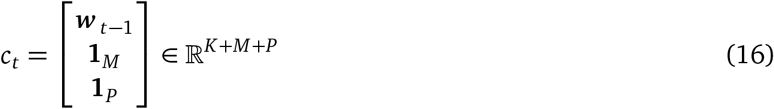

where **1**_*α*_ denotes the vector of 1’s in ℝ^*α*^ for *α* ∈ {*M, P*} (and, by convention, we set ***w*** _−1_ := 0 ∈ ℝ^*K*^). In particular, note that ***x*** _*t*_ can be recovered from the cell state via the relation 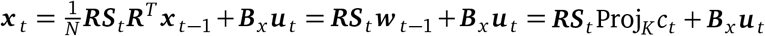, where Proj_*K*_ denotes the projection matrix that sends *c*_*t*_ onto its first *K* components, namely, ***w*** _*t*−1_.

We may also set all the parameters in *W*_*i*_ and *b*_*i*_ (part of the input gate *i*_*t*_) to be 0, so that 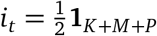(after applying the sigmoid nonlinearity). Then, define 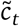 so that it equals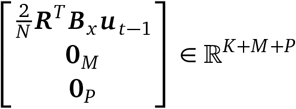. Indeed,

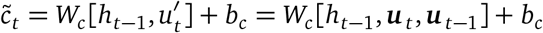

can be made to have its first *K* entries form the vector 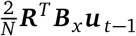 if we set *b*_*c*_ = 0 and zero out all columns of *W*_*c*_ that do not correspond to ***u*** _*t*−1_, and further by zeroing out the last *M* + *P* rows of *W*_*c*_. In block matrix form, we have defined

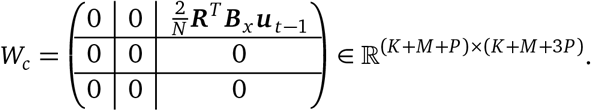

Before explicitly computing the other gates, we first analyze how the semilinear LSTM might reproduce the update for the neuromodulatory state ***z*** _*t*_ ∈ ℝ^*M*^ :

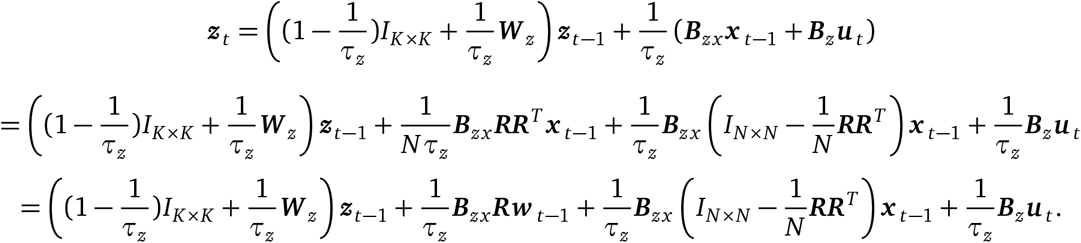

Note further that

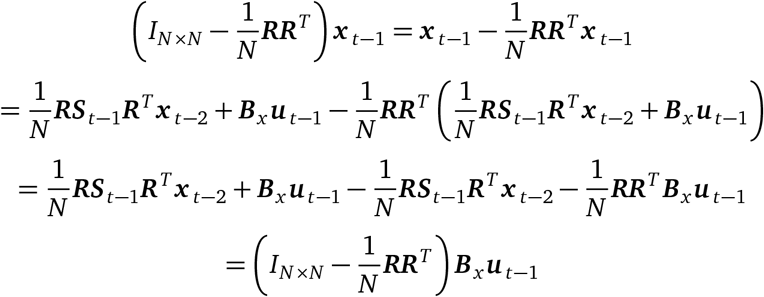

where we have again used that 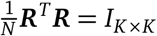. Thus, substituting this expression into our update equation for ***z*** _*t*_ above, we have

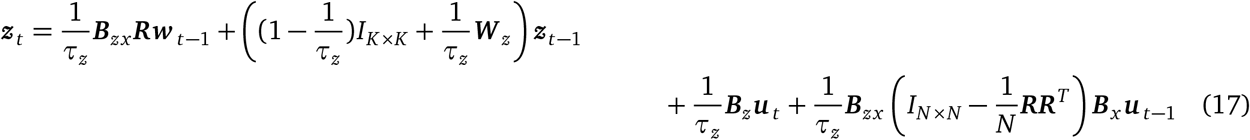

Accordingly, define the output gate *o*_*t*_ of the semilinear LSTM to be

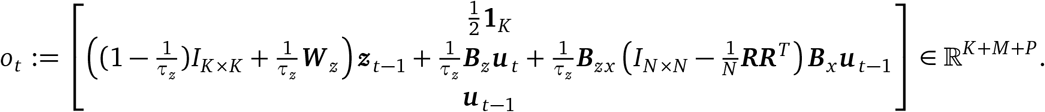

Recalling that *o*_*t*_ = *W*_*o*_[*h*_*t*−1_, ***u*** _*t*_, ***u*** _*t*−1_] + *b*_*o*_, the above gating is achieved by setting *b*_*o*_ = 0 and zeroing out the first *K* rows of *W*_*o*_. The next *M* rows of *W*_*o*_ can be set to produce the middle entry in *o*_*t*_ shown above, and the last *P* rows of *W*_*o*_ can similarly be set so as to produce the vector ***u*** _*t*−1_. That is, in block matrix form, we may take *W*_*o*_ to be

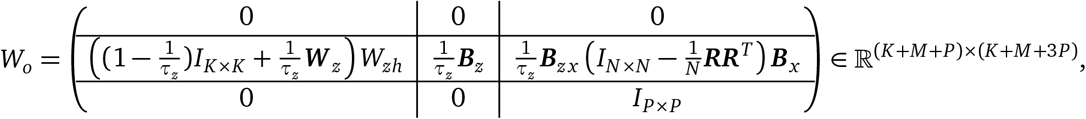

where *W*_*zh*_ ∈ ℝ^*M×*(*K*+*M*+*P*)^ is a suitable matrix (defined below) mapping the hidden state *h*_*t*_ of the semilinear LSTM to ***z*** _*t*_, so that *o*_*t*_ = *W*_*o*_ [*h*_*t*−1_, ***u*** _*t*_, ***u*** _*t*−1_]. Then, computing *h*_*t*_ = *o*_*t*_ ⊙ *c*_*t*_ ∈ ℝ^*K*+*M*+*P*^ gives

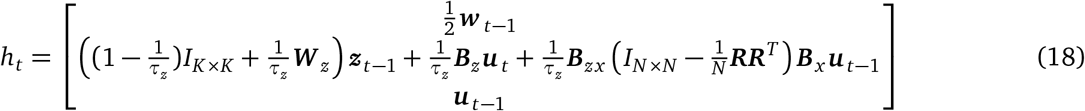

Now, observe that ***z*** _*t*_ = *W*_*zh*_*h*_*t*_, where

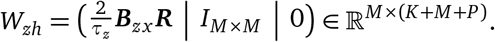

(Thus, our earlier construction of output gate *o*_*t*_ as *o*_*t*_ = *W*_*o*_[*h*_*t*−1_, ***u*** _*t*_, ***u*** _*t*−1_] is valid.) In particular, the equation ***z*** _*t*_ = *W*_*zh*_*h*_*t*_ precisely corresponds to the update equation for ***z*** _*t*_ in the linearized NM-RNN given by eq. (17). Thus, at each time *t*, the hidden state *h*_*t*_ of the semilinear LSTM is a “deconstructed” version of ***z*** _*t*_, meaning that the model effectively reproduces ***z*** _*t*_ and its dynamics through *h*_*t*_.

Finally, we set the forget gate so that it evaluates to 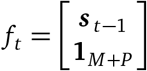. Recalling that 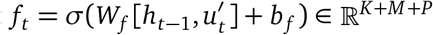, we can achieve this gating by taking the first *K* entries of *f*_*t*_ to be ***s*** _*t*−1_ = *σ*(***A***_*z*_ ***z*** _*t*−1_+***b***_*z*_) = *σ*(***A***_*z*_*W*_*zh*_*h*_*t*−1_+ ***b***_*z*_). We can also ensure that the last *M* + *P* entries of *f*_*t*_ form the vector **1**_*M*+*P*_ by zeroing out the last *M* + *P* rows of *W*_*f*_ and making the last *M* + *P* entries of the bias vector *b*_*f*_ to be sufficiently large (so that applying the sigmoid function to these entries effectively yields 1). In other words, in block matrix form, we have

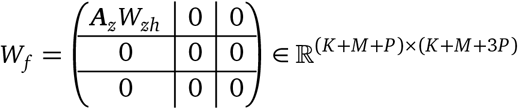

and 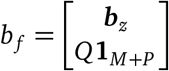 where *Q* ∈ ℝ_+_ may be chosen such that

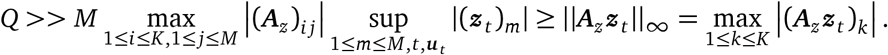

(The above supremum is taken over all components of ***z*** _*t*_, all timesteps *t* – finitely or infinitely many – and all input sequences (***u*** _*t*_)_*t*≥1_ being considered. We then invoke uniform boundedness of the components of ***z*** _*t*_ to grant the existence of such a *Q*.)

Putting together our gate computations, the cell state update equation for the semilinear LSTM (eq. (14)) can be made to reproduce the low-rank update equation of the linearized NM-RNN’s output-generating network:

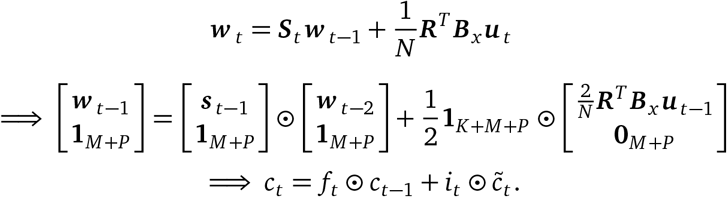

Having seen that the corresponding semilinear LSTM is capable of replicating the low-rank update equation for ***w*** _*t*_ as well as (implicitly) reproducing the update equation for ***z*** _*t*_, we at last turn to analyzing the outputs ***y*** _*t*_ generated by the NM-RNN. The output readout for the NM-RNN (at time *t* − 1, where *t* ≥ 2) can be expressed as

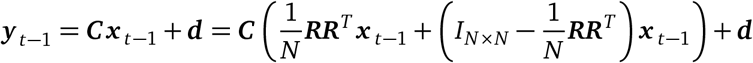

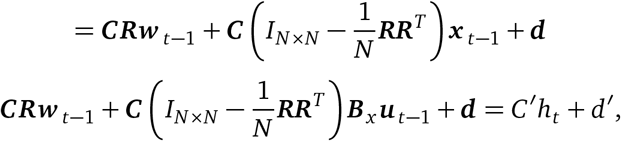

where we have used our earlier equation 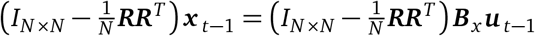. Furthermore, in the last equality, we have defined *d*^*′*^ = ***d*** and *C*^*′*^ ∈ ℝ^*O×*(*K*+*M*+*P*)^ that (in block matrix form) as

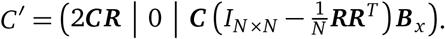

Therefore, we find that the outputs of the NM-RNN can be reproduced via a linear readout from the hidden state *h*_*t*_ of the semilinear LSTM.

Here, it should be noted that the semilinear LSTM we have constructed “lags” behind the NM-RNN by one timestep, i.e., the hidden state *h*_*t*_ is used to produce the (*t* − 1)th output state ***y*** _*t*−1_. This is a consequence of the fact that at each timestep in the NM-RNN, the neuromodulatory state ***z*** _*t*_ is updated before ***x*** _*t*_, whereas in the semilinear LSTM, the cell state *c*_*t*_ (loosely corresponding to ***x*** _*t*_) is updated before the hidden state *h*_*t*_ (loosely corresponding to ***z*** _*t*_); as a result, the outputs of the semilinear LSTM are staggered by one timestep. In practice, this does not change the fact that the semilinear LSTM can replicate the outputs of the NM-RNN.

Having constructed a semilinear LSTM with hidden size *K* + *M* + *P* that reproduces the states of the linearized NM-RNN, we finally turn to counting the total number of learnable parameters used by the semilinear LSTM (along with its linear readout):

1. *f*_*t*_ : *W*_*f*_ only transforms the hidden state *h*_*t*_ (i.e., 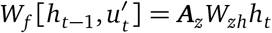, where ***A***_*z*_*W*_*zh*_ ∈ ℝ^*K×*(*K*+*M*+*P*)^), giving us *K*(*K* + *M* + *P*) parameters. The bias vector *b*_*f*_ gives an additional *K* + *M* + *P* parameters, for a total of (*K* + 1)(*K* + *M* + *P*) parameters.
2. *i*_*t*_ : We zero out all of these parameters, giving us a count of 0.
3. 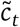 : The only parameters used in this computation are those that linearly transform ***u*** _*t*−1_ into *K*-dimensional space (i.e., the elements of 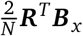), so the parameter count here is *PK*.
4. *o*_*t*_ : The bias term *b*_*o*_ was set to 0 and we defined the matrix *W*_*o*_ so that the middle *M* rows of *W*_*o*_ contained nontrivial entries, and the bottom *P* rows have *P* nontrivial parameters stemming from the identity matrix *I*_*P×P*_. Consequently, we obtain a contribution of *M* (*K* + *M* + 3*P*) + *P* parameters from the output gate computation.
5. Readout: We have that *d*^*′*^ ∈ ℝ^*O*^ and *C*^*′*^ ∈ ℝ^*O×*(*K*+*M*+*P*)^ uses a total of *O*(*K* + *P*) nontrivial parameters, giving us a total of *O*(*K* + 1 + *P*) parameters used.

Thus, the number of nontrivial parameters used by this semilinear LSTM is

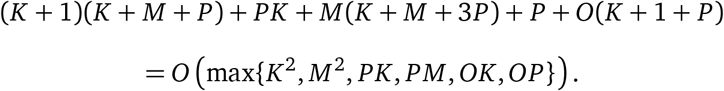

## B Derivation of exact neuromodulatory signal for rank-1 NM-RNNs

In this section, we precisely quantify how the neuromodulatory signal in a rank-1 NM-RNN is constrained by the target output signal. First, we derive the exact neuromodulatory signal in a general (possibly nonlinear) rank-1 NM-RNN that has successfully learned to produce a target output signal *f* (*t*) ∈ ℝ. Then, we specialize to the case in which the rank-1 NM-RNN has linear dynamics.

We start with a general rank-1 NM-RNN that produces the scalar output ***y*** (*t*) ∈ ℝ at each time *t*, and for which the output-generating network’s nonlinearity is denoted by *ϕ* (which could be tanh, but also any other function). Because *K* = 1, we have ***L*** ∈ ℝ^*n×*1^, ***R*** ∈ ℝ^*n×*1^. Furthermore, treating ***L*** and ***R*** as vectors in ℝ^*n*^, let 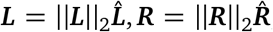, where 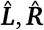 are unit vectors, and define 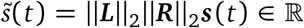. Additionally, let ***u***(*t*) ∈ ℝ^*P*^ denote the input signal over time. Then, for *t >* 0, the dynamics of the output-generating network read:

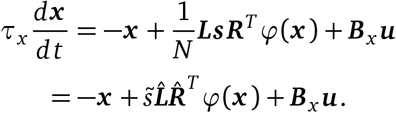

we now define t he r ank-1 d ynamics v ariable 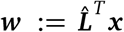 and a residual mode 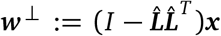, so that 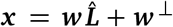 is a combination of a rank-1 mode and a residual component. This gives us the dynamics equations

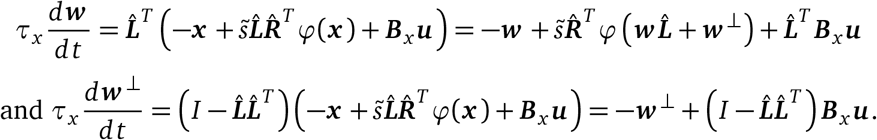

Using the basic theory of ordinary differential equations, we may solve the latter ODE to obtain

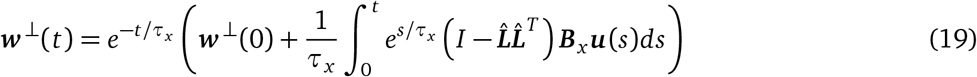

As a special case, in the absence of any inputs, we would have 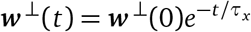. For ease of notation, we define the function *J* : ℝ → ℝ^*N*^ by

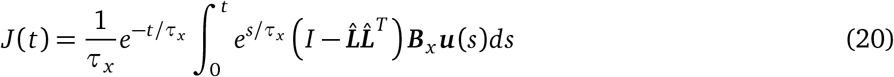

Now, recall that our desired output signal is some prespecified function *f* (*t*). Letting the linear readout (of the output-generating network) be given as 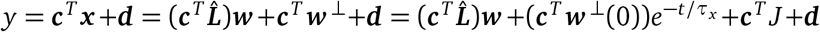, then solving for ***w*** and differentiating yields the equations

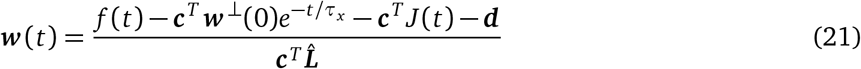

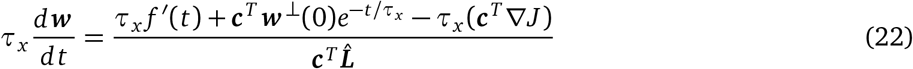

Now, observe that

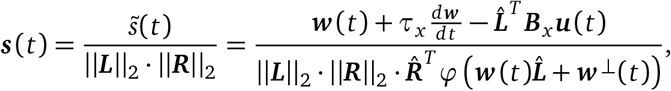

meaning that

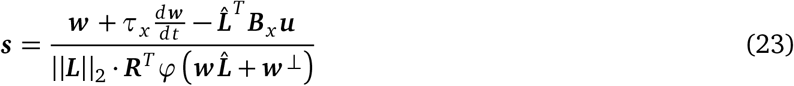

Substituting in our earlier expressions for ***w*** (*t*) and 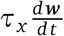 into the formula for ***s*** (*t*), we obtain the formula

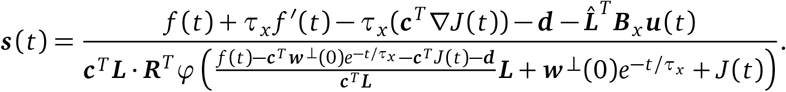

Moreover, in the absence of any inputs to the system, the neuromodulatory signal would be

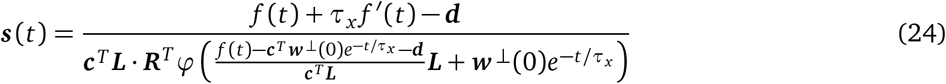

Furthermore, if we also know that that *τ*_*x*_ is sufficiently small and that ***w*** ^⊥^(0) has sufficiently small entries, then (owing to the exponential decay of ***w*** ^⊥^) we may further approximate ***s*** as

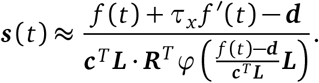

Thus, for NM-RNNs in which the output-generating network’s nonlinearity is removed (i.e., where *ϕ*(***x***) := ***x***), eq. (24) implies that, in the absence of inputs, the neuromodulatory signal is

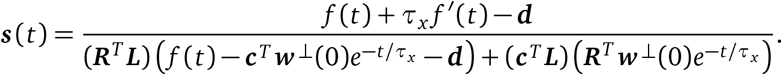

If we assume further that ***w*** ^⊥^(0) is sufficiently small and *τ*_*x*_ is also sufficiently small, then we may make the approximation

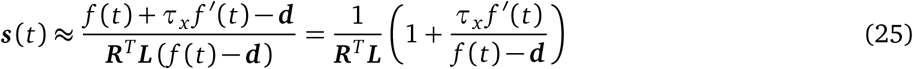

In particular, for *f* (*t*) a linear ramp (such as during the ramping phase of the MWG task), eq. (25) implies that ***s*** (*t*) should follow a power law.

## C Computing details

### C.1 Code, data, and instructions

The code required to reproduce our main results is included in the nm-rnn GitHub repository: https://github.com/lindermanlab/nm-rnn. All code was written using Python, using fast compilation and optimization code from the JAX and Optax packages [40, 41]. Experiments and models were logged using the Weights and Biases ecosystem [42].

#### Instructions

To generate the results that we have presented, we have included the package folder “nmrnn” and folder of training scripts “scripts”. The packages needed to replicate our coding environment are in “requirements.txt”.

Each training script has a “config” dictionary near the top of the file that allows you to set hyperparameters, it is specifically set up to work with Weights and Biases. In the Jax framework, different initializations are set by changing the “keyind” value in the config dictionary.

### C.2 Data generation

All data was generated synthetically.

#### Rank-1 Measure-Wait-Go

Rank-1 linearized NM-RNNs were trained on 40 trials, generated using the target intervals [12, 14, 16, 18] and by setting the (integer-valued) timestep at which the measure cue appeared to be between 10 and 19, inclusive. The delay period was fixed to be 15 (timesteps). For any given target interval *T*, the target output ramp was the ramping function *f*_*T*_ (*t*) given by

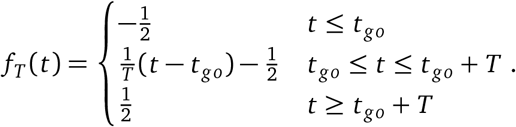

The total number of timesteps was set to be 110. In generating Figure 3B of the main text, a trained network was tested on the intervals 7 (extrapolation below), 15 (interpolation of trained intervals), and 23 (extrapolation above). The theoretically expected neuromodulatory signal ***s*** (*t*) and rank-1 state *h*(*t*) were computed using eqs. (8) and (21), respectively.

#### Rank-3 Measure-Wait-Go

All networks were trained on 40 trials, generated using desired intervals [12, 14, 16, 18] and by setting the integer-valued delay period (interval between wait and go cues) between 10 and 19. Networks were tested on these trials plus corresponding extrapolated trials with interval lengths [4, 6, 8, 10, 20, 22, 24, 26].

#### Rank-3 Multitask

Initial training was completed on 3000 randomly sampled trials from [DelayPro, DelayAnti, MemoryPro], with random angles and task period lengths. Retraining was completed on 1000 trials of MemoryAnti with random angles and task period lengths. Testing was completed on a different, fixed set of 1000 samples of each task.

#### Element Finder Task

All networks were trained on one-dimensional input sequences of length 26. For each input sequence used during training, the input at the first timestep was the query index *q*, which was uniformly sampled at random from {0, 1, …, 24}. Each input in the proceeding sequence of 25 inputs was uniformly sampled at random from {−10, −9, …, 9, 10}.

### C.3 Training details

Training was performed via Adam with weight decay regularization and gradient clipping [43, 44]. Specific AdamW hyperparameters varied by task, see below for exact details.

#### Rank-1 Measure-Wait-Go

Training was performed via full-batch gradient descent for 50k iterations using the standard Adam optimizer with learning rate 10^−3^. The linearized NM-RNN trained had the hyperparameters *N* = 100, *M* = 20, *K* = 1, *τ*_*x*_ = 2, *τ*_*z*_ = 10. Training took about 20 minutes. A single network was trained to produce the results shown in Figure 3B, and many more networks were trained while producing preliminary results.

#### Rank-3 Measure-Wait-Go

Training was performed with initial learning rate 10^−2^. For NM-RNNs, we first trained the neuromodulatory subnetwork parameters and output-generating subnetwork parameters separately for 10k iterations each, followed by training all parameters for 50k iterations. We set specific hyperparameters as follows:

- **NM-RNN:** *N* = 100, *M* = 5, *K* = 3, *τ*_*x*_ = 10, *τ*_*z*_ = 100.
- **LR-RNN:** As above, except *N* = 106 for parameter matching.

Models were trained using at least 32 parallel CPUs (1G memory each) on a compute cluster. Training the NM-RNNs took about 15 minutes each, and training the LR-RNNs took about 9 minutes each. Ten of each model were used to produce the results shown in the paper; however, we trained many more while producing preliminary results.

#### Rank-3 Multitask

Training on first three tasks was performed with initial learning rate 10^−3^, for 150k iterations. Retraining on the MemoryAnti task was performed with initial learning rate 10^−2^ for 50k iterations. We set specific hyperparameters as follows:

- **NM-RNN:** *N* = 100, *M* = 20, *K* = 3, *τ*_*x*_ = 10, *τ*_*z*_ = 100.
- **LR-RNN:** As above. In this case, since the LR-RNN receives contextual inputs as well as sensory/fixation inputs (compared to the NM-RNN’s output generation subnetwork which only receives sensory/fixation inputs), the LR-RNN has more parameters.

Models were trained using at least 32 parallel CPUs (1G memory each) on a compute cluster. Training the NM-RNNs took about 2 hours each, and training the LR-RNNs took about 2.5 hours each. These times include both training and retraining. Ten of each model were used to produce the results shown in the paper; however, we trained many more while producing preliminary results.

#### Element Finder Task

For all networks, training was done over 20k iterations of gradient descent using a batch size of 128. Each batch consisted of newly randomly generated input sequences. The standard Adam optimizer was used, and the learning rate and hyperparameters were varied across the different models tested:

- **LSTM:** We trained LSTMs of hidden size *N* = 10 using a learning rate of 10^−2^.
- **NM-RNN:** We trained multiple NM-RNNs across the hyperparameter combinations (*M, N, K*) = {(5, 18, 8), (5, 13, 12), (10, 14, 5), (10, 12, 7), (15, 6, 5)}, fixing *τ*_*x*_ = 10 and *τ*_*z*_ = 2, and usinga learning rate of 10^−2^.
- **LR-RNN:** We trained multiple LR-RNNs across the hyperparameter combinations (*N, K*) = {(23, 10), (31, 7), (50, 4), (83, 2)}, fixing *τ*_*x*_ = 10, and using a learning rate of 10^−2^.
- **(Full-rank) RNN:** We trained multiple RNNs with hidden size *N* = 16, fixing *τ*_*x*_ = 10, and across the learning rates {10^−3^, 10^−2^, 10^−1^}.

All trained models were parameter-matched (∼ 500 total parameters). For each model type, hyperparameter combination, and learning rate, ten such models were trained; the resulting model performances are illustrated in Figure 5B of the main text. Figure 5C was illustrated using a single run of an LSTM (*N* = 10), a full-rank RNN (*N* = 16), and three NM-RNNs ((*M, N, K*) = (5, 18, 8), (5, 13, 12), (10, 12, 7)), where all models used a learning rate of 10^−2^. Finally, Figure 5D-F show results for a single NM-RNN ((*M, N, K*) = (5, 18, 8), lr = 10^−2^). Training each network took roughly 5 minutes. Many more models were trained while producing preliminary results.

### C.4 Metric for multitask setting

For the multitask setting, we used the percent correct metric from [1]. A trial was counted as correct if (1) the fixation output stayed above 0.5 until the fixation input switched off, (2) the angle read out at the final timestep was within *π/*10 of the desired angle.

## Supplemental figures

**Figure S1:**
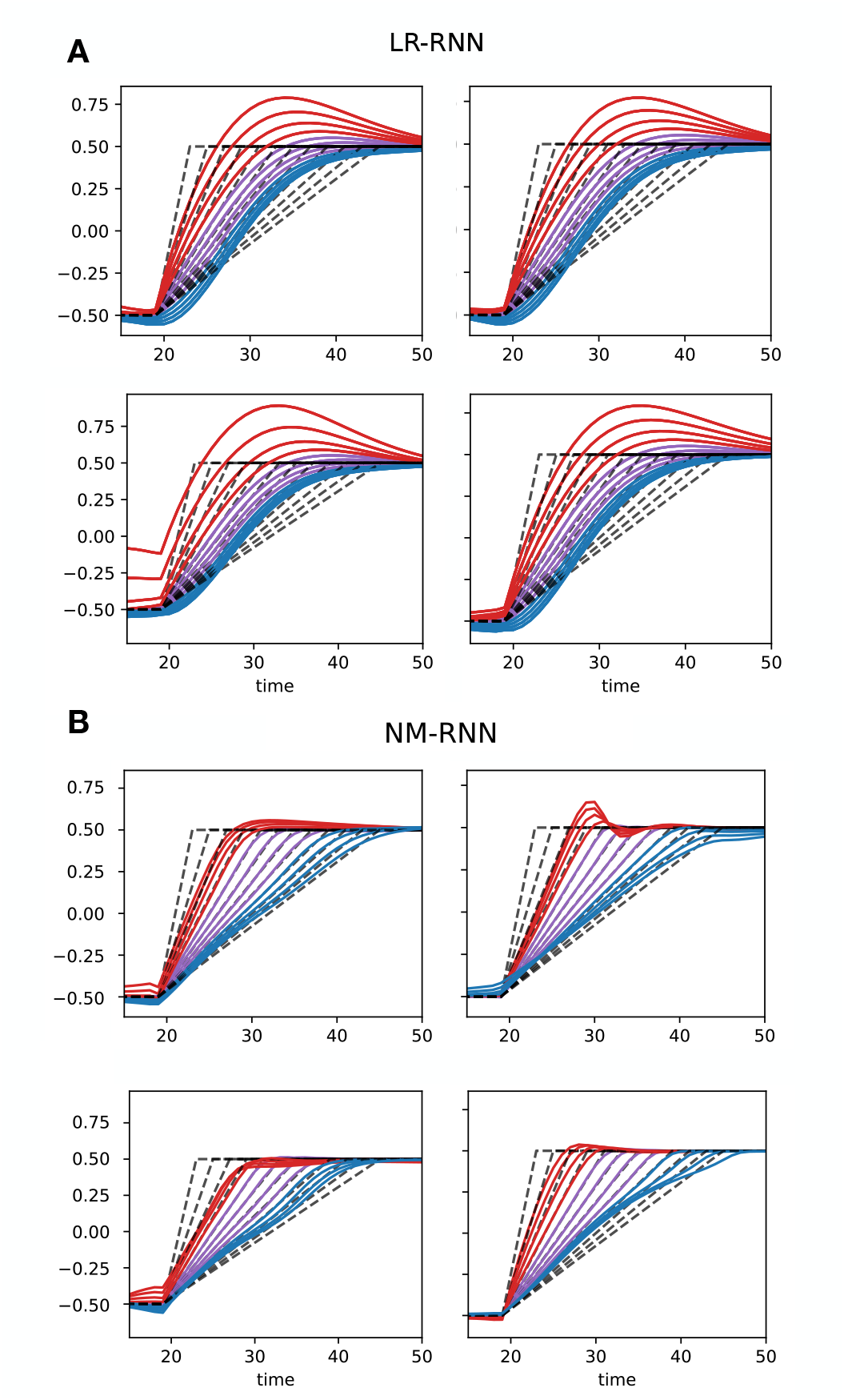
Example output comparison plots for the measure-wait-go task, as in fig. 3D in the main text, for four additional trained (**A**) low-rank RNNs and (**B**) NM-RNNs. Colors indicate extrapolated/trained intervals as in the main text.

**Figure S2:**
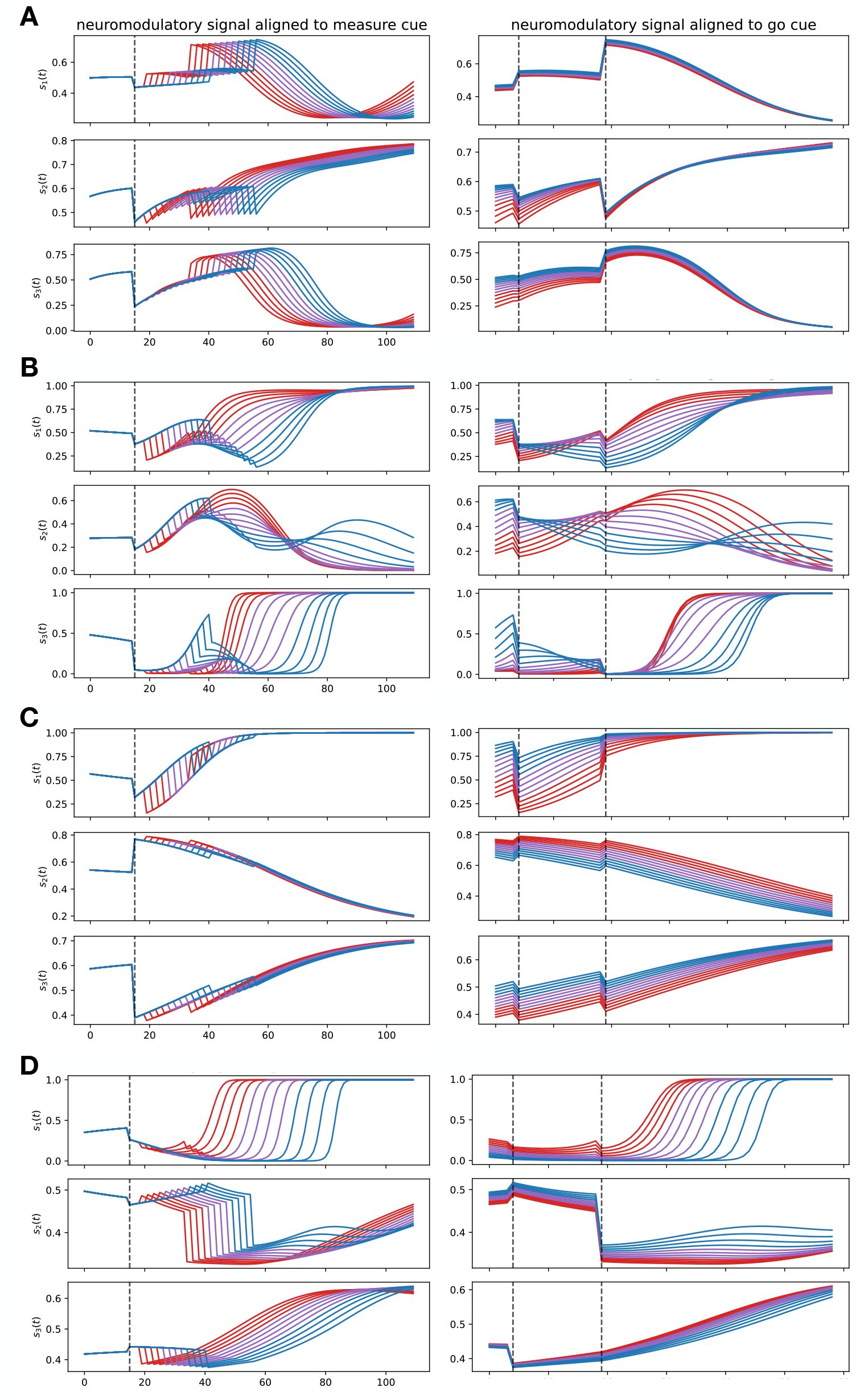
Example neuromodulatory signal plots for the measure-wait-go task, as in fig. 3E in the main text, for four additional trained NM-RNNs (same networks as shown in fig. S1B). Colors indicate extrapolated/trained intervals as in the main text.

**Figure S3:**
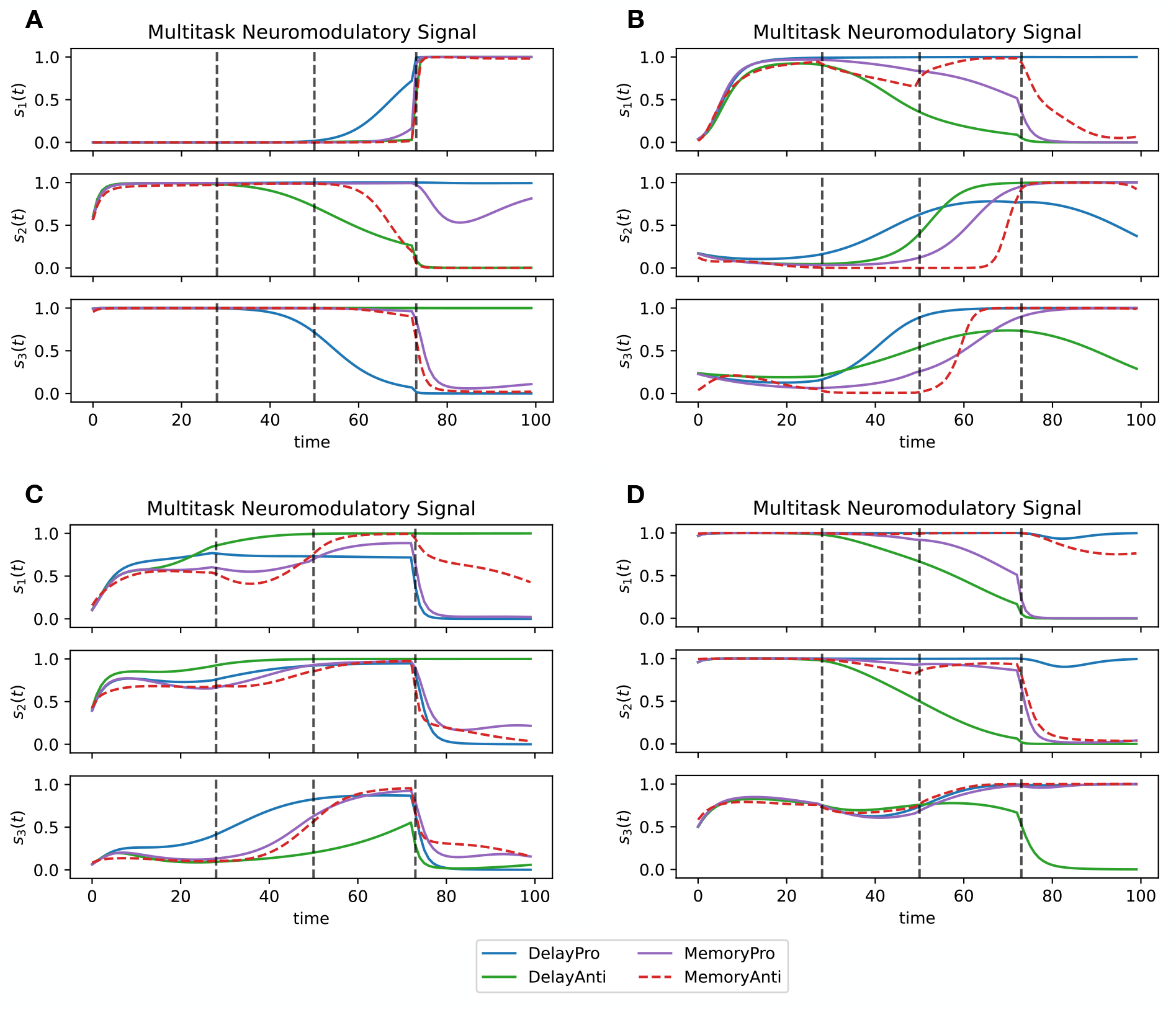
Example neuromodulatory signal plots for the multitask setting, as in fig. 4C in the main text, for four additional trained networks.

**Figure S4:**
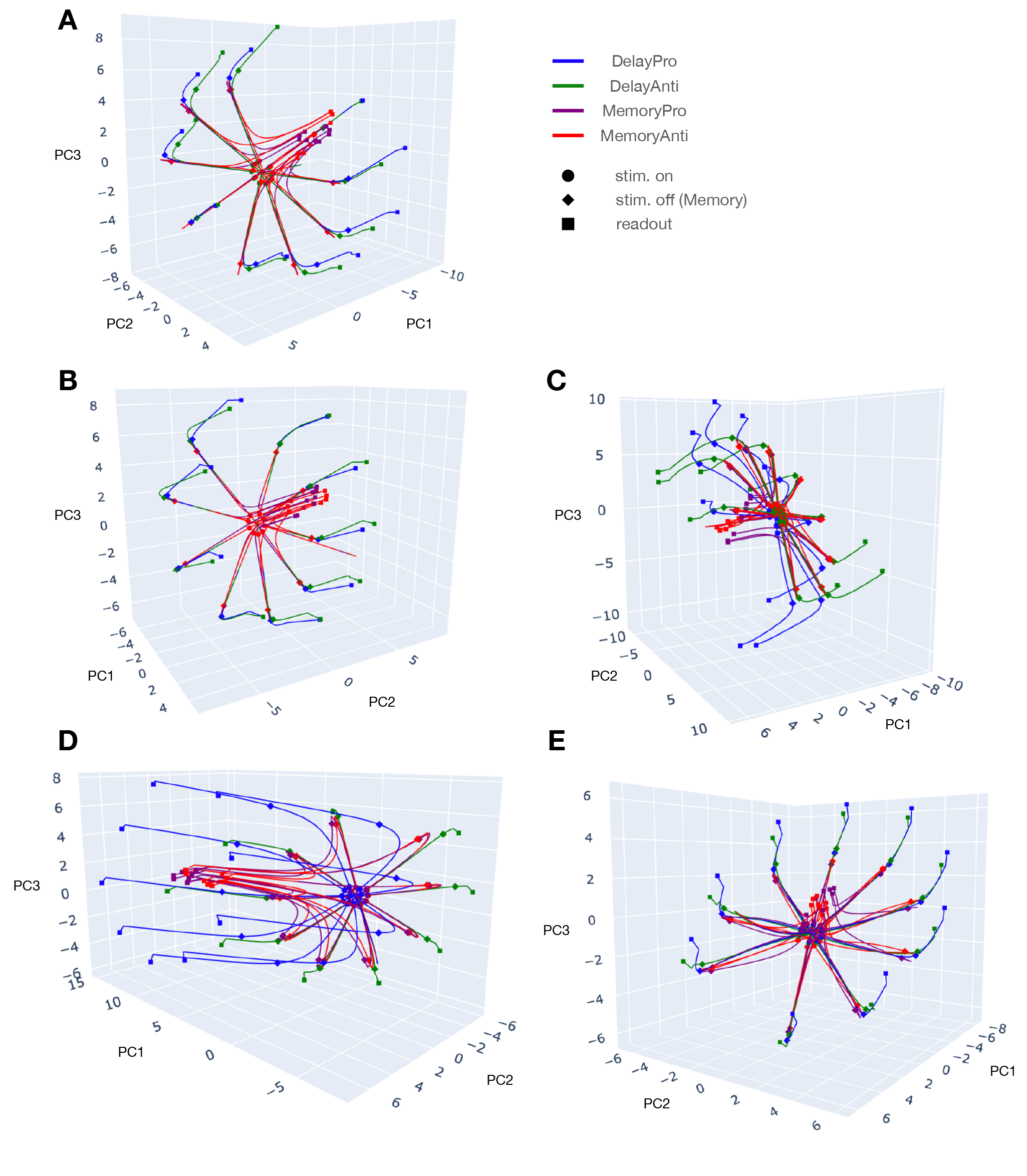
First three PCs of neural activity in multitask setting, plotted until readout period (for ease of visualization). **A**. Network visualized in main text, **B-E**. Four additional networks.

**Figure S5:**
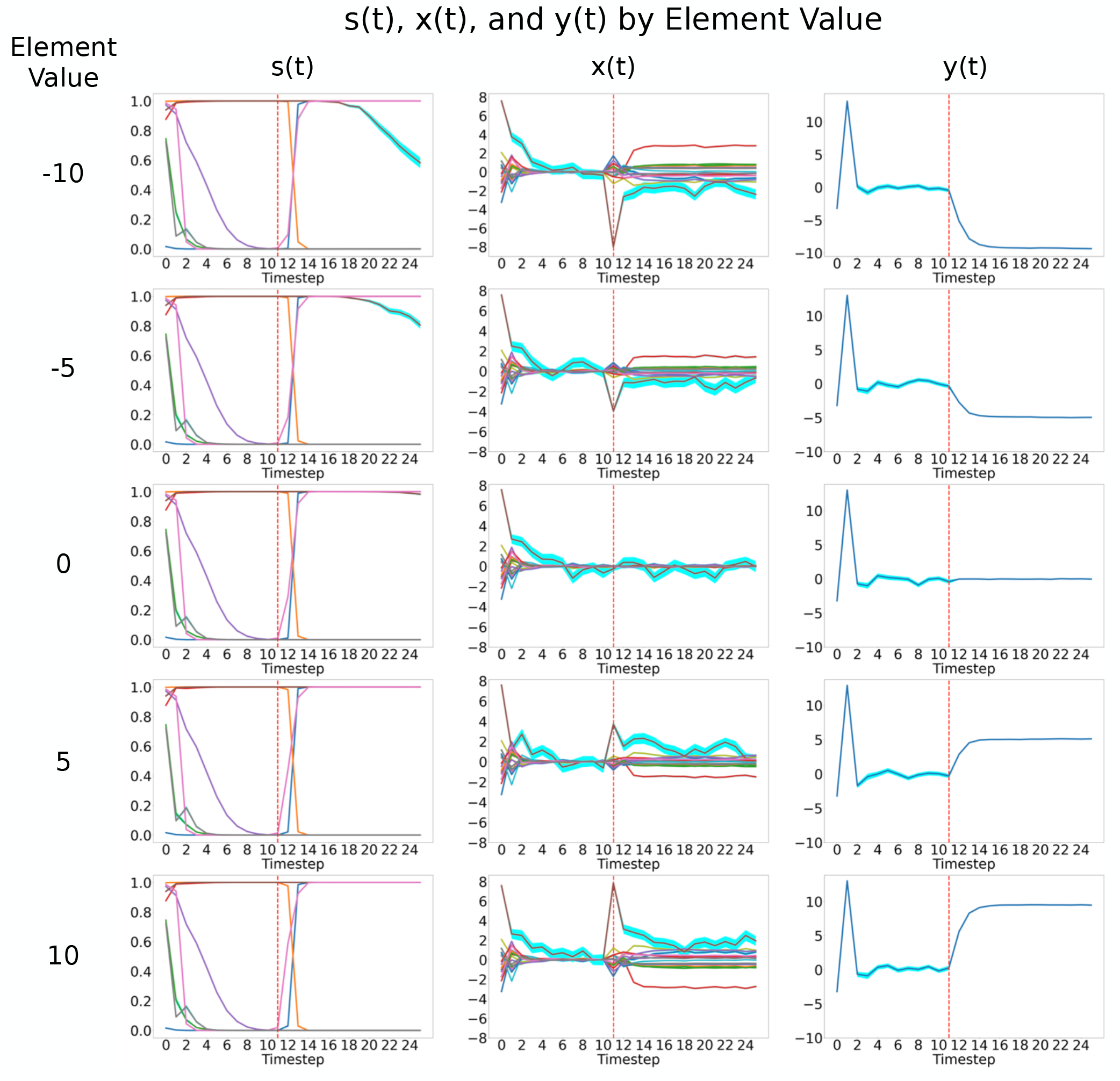
Sample internal states of an NM-RNN (*M* = 5, *N* = 18, *R* = 8) trained on the Element Finder Task, shown for 5 different element values (−10, −5, 0, 5, and 10). Each plot shows how all of the components of one of the vectors *s*(*t*) (left), *x* (*t*) (middle), *y*(*t*) (right) vary through time. The query index is fixed to be 10, as indicated by the red dashed line in each plot. Each line shown is averaged over 100 independent runs of the model (standard error shown in cyan).

**Figure S6:**
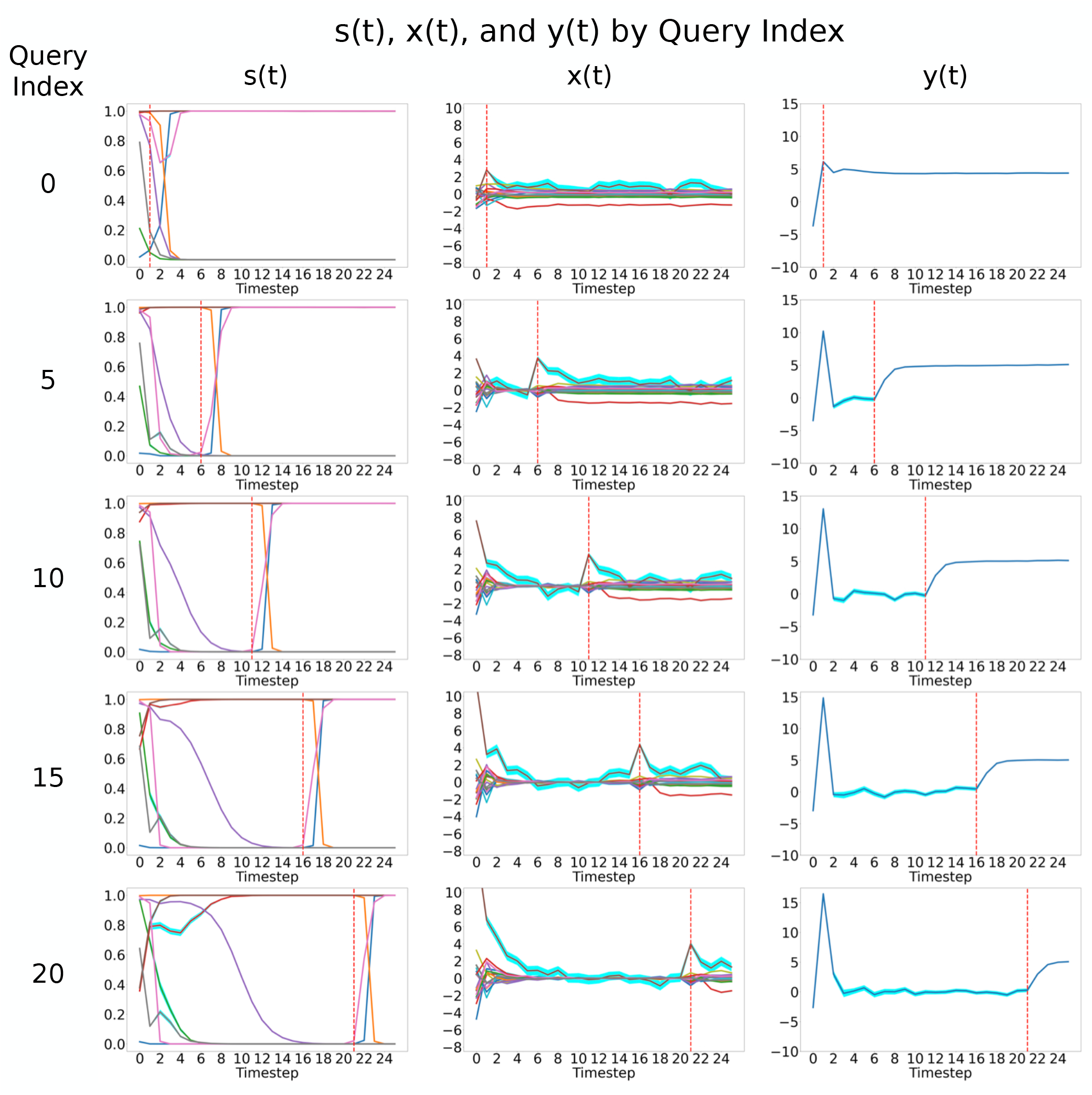
Sample internal states of an NM-RNN (*M* = 5, *N* = 18, *R* = 8) trained on the Element Finder Task, shown for 5 different query indices (0, 5, 10, 15, and 20), while fixing the target element value to be 5 in each case. Each plot shows how all of the components of one of the vectors *s*(*t*) (left), *x* (*t*) (middle), *y*(*t*) (right) vary through time. In each plot, the onset of the query index is indicated by the red dashed line. Each line shown is averaged over 100 independent runs of the model (standard error shown in cyan).

